# Lead exposure positively affects coccidia abundance in feral pigeons (*Columba livia*)

**DOI:** 10.1101/2025.02.25.640109

**Authors:** Aurélie Jeantet, Fabienne Audebert, Simon Agostini, Beatriz Decencière, Lou Gimeno, Tom Compain, Pierre Federici, Camille Lorang, David Rozen-Rechels, Julien Gasparini

## Abstract

1. In urban environments, organisms are exposed to high levels of pollutants, including trace metals, whose concentrations can be increased by anthropogenic activities. Numerous studies have shown the toxic effects of pollutants on exposed organisms. However, due to their life within the host, parasites can also be affected by exposure to these pollutants. Overall, previous correlative findings reveal that the effects of parasite exposure to pollutants through the host can vary, from positive to negative, highlighting the complexity of host-parasite-environment interactions in relation to pollution.
2. In this study, we experimentally tested whether lead exposure affects the abundance of ecto-, meso-, and endoparasites in wild pigeons (*Columba livia*), which host a wide variety of parasites and are naturally exposed to trace metals in urban environments. As we had previously reported the toxic effects of lead on the immune system of pigeons, we expected lead exposure to be indirectly beneficial for the parasites.
3. To test this, we used a sample of wild pigeons captured in Paris, half of which were experimentally exposed to lead concentrations similar to those found in Paris. We then measured the abundance of several parasites: blood parasites (hemosporidian parasites), ectoparasites (*Columbicola columbae* and *Campanulote compar*), coccidia and helminths. Additionally, we measured the intensity of the pigeons’ antiparasitic behavioral response (grooming) through behavioral analyses.
4. Our results did not reveal toxic effects of lead exposure on the parasites. On the contrary, we found positive effects of this exposure on coccidia abundance in male pigeons. This result could be explained by a toxic effect of lead on the host’s antiparasitic immune strategy, making the hosts less hostile to parasites. However, we found no effect of lead exposure on the abundance of lice, blood parasites and helminths. This could be explained by the lack of impact of lead exposure on grooming antiparasitic activity in our experiment, and by seasonal variations in the activity of blood parasite vectors, which may have masked potential effects of our treatment.
5. In conclusion, our study highlights the importance of considering the host environment, particularly pollutants, to understand parasite dynamics.

## Introduction

Parasites are ubiquitous and induce high pressure on living organisms (Wobeser, 2008). Indeed, according to Price (1980), any living organism is affected by parasitism, as a host or as a parasite. Thus, the parasitic lifestyle concerns more than half of living species and 50% of plants and animals are parasitized during at least one stage of their life cycle (Tirard and al, 2016). Parasitism is defined as a sustainable ecological interaction between individuals of two species in which a parasitic organism draws its resources from another organism called host, decreasing its fitness (Wobeser, 2008). In response to pressures from parasites, anti-parasitic strategies have been favored by selection, and induce pressures on parasites through behavioral and immune responses (Poirotte, 2016). Thus, in addition to the use of its resources, the host can suffer from tissue damage caused by parasites or its own defenses. Indeed, parasites can be responsible for accumulations of energy costs over time due, for example, to energy losses linked to the immune response (Koop et al., 2011; Wobeser, 2008). Consequently, the host could suffer of a reduction in competitive abilities or an increase in sensitivity to other stress factors when infected (Wobeser, 2008).

In addition to the cost induced by parasites, hosts can suffer harmful effects from other environmental factors, such the pollutant exposure, elevated temperature or salinity. These environmental effects could influence the presence of parasites and their effects on hosts (Lewis et al., 2003; Marcogliese, 2008). Indeed, due to their lifestyle within the host, parasites are subject to the same environmental conditions (harmful or not), which can also impair cost on parasite. Therefore, host exposure to pollutants may be profitable or not for the parasites (Eeva et al., 2005).

First, parasites exposed to pollutants through the host may suffer directly from the toxic effects of this exposure. For example, Lefcort et al. (2002) showed negative effects of pollutants on the intensity and diversity of parasites in two host species of snails (*Physella columbiana* and *Lymnaea palustris*). Similarly, El-Bouhy et al. (2016) also reported negative effects of pollutants on fish ectoparasites (*Oreochromis niloticus*). Second, host exposed to pollutants may induce an increase of parasites, which could be explained by toxic effects of pollutants on host immunity (Bagge & Valtonen, 1996; Boyce & Yamada, 1977; Pascoe & Cram, 1977). Accordingly, Sanchez-Ramirez et al. (2007), observed that exposure to polluted sediments increased the abundance of ectoparasites *Cichlidogyrus sclerosus* in fish (Nile tilapia, Oreochromis niloticus). Also, Korine et al. (2017) showed that the abundance of bat ectoparasites (*Pipistrellus kuhlii*) was significantly higher when bats foraged over polluted water. Finally, Gasparini et al. (2014) observed a positive correlation between the concentration of lead in the feathers of Parisian pigeons and the intensity of haemosporidian parasites.

Overall, these previous correlative results reveal that the effects of pollutants exposure of parasites through the host can vary, highlighting the complexity of host-parasite-environment interactions in relation to pollution (El-Bouhy et al., 2016). However, experimental studies are now required to confirm these previous results.

In this study, we experimentally tested whether lead exposure affects positively or negatively the abundances of ecto, meso and endo-parasites in feral pigeons (*Columba livia*). Feral pigeon represents a good model to test our hypotheses because birds host a wide diversity of parasites including ectoparasites, which live outside the host (e.g. lices), mesoparasites which live inside the body of the host, with access to the external environment (e.g. intestinal parasites) and endoparasites, which live inside the body of the host, without access to the external environment (e.g. blood parasites). In addition, pigeons mainly exploit urban environments and are therefore naturally exposed to trace metals as lead (Chatelain et al., 2014). As we previously reported toxic effects of lead on the immune system of pigeons (Chatelain, Gasparini, & Frantz, 2016a, 2016b; Jeantet et al., 2023), we expected that lead exposure can indirectly be profitable for endo- and mesoparasites. As for ectoparasites the pigeon’s antiparasitic defenses is mainly the grooming, we also experimentally tested whether lead exposure impaired the grooming. In case of negative effect of lead exposure on grooming, we would expect a positive effect of lead exposure on ectoparasites abundances.

To test these predictions, we used parasites data collected in parallel of a previous experiment testing another hypothesis on the half of pigeons non-exposed to the anthelmintic treatment (Jeantet et al., 2024). In this context, we used 66 feral pigeons captured in Paris, France and housed in aviaries at the CEREEP (Centre for Research in Experimental and Predictive Ecology) biological station. Half of these 66 pigeons were exposed to lead for a period of 6 months and we measured the abundance of several parasites: blood parasites (*haemosporidian parasites*), ectoparasites (*Columbicola columbae* and *campanulote compar*), coccidia and helminths. In addition, we measured the intensity of the behavioral anti-parasite response (grooming), through behavioral scans.

## Material and methods

For this study, we used a subsample of pigeons (n = 66) non-exposed to the anthelmintic treatment of a previous study testing another hypothesis. The sampling and the general protocol of lead exposure are reported in Jeantet et al. (2024). Hereafter, we briefly summarize the bird sampling and the lead exposure protocol.

### Bird sampling

In January 2022, a sample of 66 pigeons was caught in Paris by Julien Gasparini, Aurélie Jeantet, Fabienne Audebert and David Rozen-Rechels. Then in order to identify them individually, we put a numbered and colored ring on their left paw and a colored ring on their right paw. They were distributed in six outdoor aviaries (dimensions: 3 m x 2,2 m x 2,2 m) in a way to equilibrate body mass (ANOVA, F_5, 60_ = 0.15, P-value = 0.98), melanin-based plumage coloration (Kruskall-Wallis, χ^2^_5_ =0.62, P-value = 0.99), sex (GLM, χ^2^_5_ =2.27, P-value = 0.81) and site of capture (Fisher’s exact test, P-value = 0.99) at the CEREEP (Centre de Recherche en Écologie Expérimentale et Prédictive, UMS 3194, 48° 17′ N, 2° 41′ E).

### Lead exposure

All birds were placed in the aviaries at least two weeks before the start of the experiment for acclimation. A lead-exposure treatment was assigned to birds of the six aviaries, with two experimental groups and three replicates per group: lead-exposed pigeons (1st experimental group: n=33) and non-lead-exposed pigeons (2nd group: n=33). For lead-exposed pigeon, we placed lead acetate diluted in tap water into drinking troughs and baths. The lead concentration used was 10 ppm lead acetate, based on Parisian lead pollution (Chatelain, Gasparini, & Frantz, 2016a; Chatelain, Gasparini, Haussy, et al., 2016). For control pigeons (non-lead-exposed pigeons), drinking troughs and baths were filled with tap water only. During all the experiment, birds were fed ad libitum with a mix of maize, wheat, and peas, and aviaries were enriched with baths and branches as perches. At the end of the experiment, all birds were released into the wild near their site of capture.

### Lice abundance (ectoparasite abundance)

We quantified the abundance of two species of lice *Columbicola columbae* and *Companulotes compar*, we ran visual examinations following the method described in Koop & Clayton (2013).

The counts were realized on weeks 10 and 24. Each pigeon was observed five minutes through a standardized protocol, first the right wing and then the left wing were examined for a duration of 1’30 per wing, following by the examination of the neck, de head and the body also for 1’30 and at last, the tail was observed for a duration of 30 seconds. All birds were observed by AJ, and each part of the body was observed once to avoid counting many time the same lice. The variable used to quantify lice abundance was the total number of lice of both species counted during this standardized 5-minute protocol for each bird.

### Haemosporidian parasite abundance (endoparasite abundance)

To estimate the abundance of Haemosporidian parasites (*Heamoproteus* spp., *Plasmodium* spp. and *Leucocytozoon* spp.), we used a method adapted from (Jacquin et al., 2011). Briefly, blood smears were realized just before the start of the experimental treatment and every month throughout the experiment, i.e. seven samples for each pigeon (t0 to t6 sessions). All smears were fixed in methanol and stained by using May-Grünwald Giemsa staining (Sordolab kit, ref COLRASA) to reveal Haemosporidian parasites. The smears were then photographed using an optical microscope equipped with a camera and the blood parasites were enumerated by counting the number of infected blood cells out of 5000 blood cells counted, using ImageJ software.

### Coccidia and helminth abundances (endo- and mesoparasite abundances)

We indirectly quantified coccidia and helminths in the feces, using a coproscopy method. Coproscopies were realized using Macmaster flotation technique, feces of each pigeon were collected then weighed and floated in a Falcon tube of 15 mL with a solution of saturated NaCl. After shaking the tubes and waiting for 15 minutes, the supernatant was spread on a Macmaster slide and coccidia and helminth eggs were counted using an optical microscope (modified from Raynaud et al. (1970) method).

### Preening activity

In order to quantitatively measure preening activity, we ran behavioral observations during 9 weeks from the 6 to the 24. The pigeons were each observed for 30 minutes during weeks 6 and 8 (15 minutes in the morning and 15 minutes in the afternoon) and 15 minutes during weeks 7, 15, 16, 18, 20, 22 and 24, bringing the total observation time of each pigeon at 165 minutes. During the scans, observers were placed equidistant from the aviaries (Fig. 5a). To limit bias among observers and time, the order in which the aviaries were observed as well as the observers varied between each scan. Furthermore, the same observer has never observed the same pigeon twice in a row. Every minute the observers noted the behavior that the pigeons assigned to them performed and among all the observed behaviors. The variable used to quantify the grooming was, therefore, the total number of times the pigeons were observed displaying a preening behaviors during the 165 minutes of observation.

### Statistics

Statistical analyses were performed using R (version 4.1.2). To test the effect of lead exposure on lice, coccidia, helminths and blood parasites abundances and on the preening behavior, we ran five generalized linear mixed models (GLMMs, one model for each variable) with a negative binomial distribution (glm.nb function from the MASS package). Sex and aviary were added as a cofactor for the four models, as a fixed factor and a random factor, respectively. For lice and blood parasites abundances, the measurements were performed at different times, therefore we added the time (weeks for lice abundances and the month session t0 to t6 for blood parasites) as a cofactor and the pigeon ID nested in the aviary as a random factor. Also, for the blood parasite abundance, we added the initial blood parasite abundance as a covariate. To test the effect of lead exposure on coccidia abundance, we ran the model with and without an extreme value of our dataset in the non-lead-exposed female group (Fig. 1).

**Figure 1.**
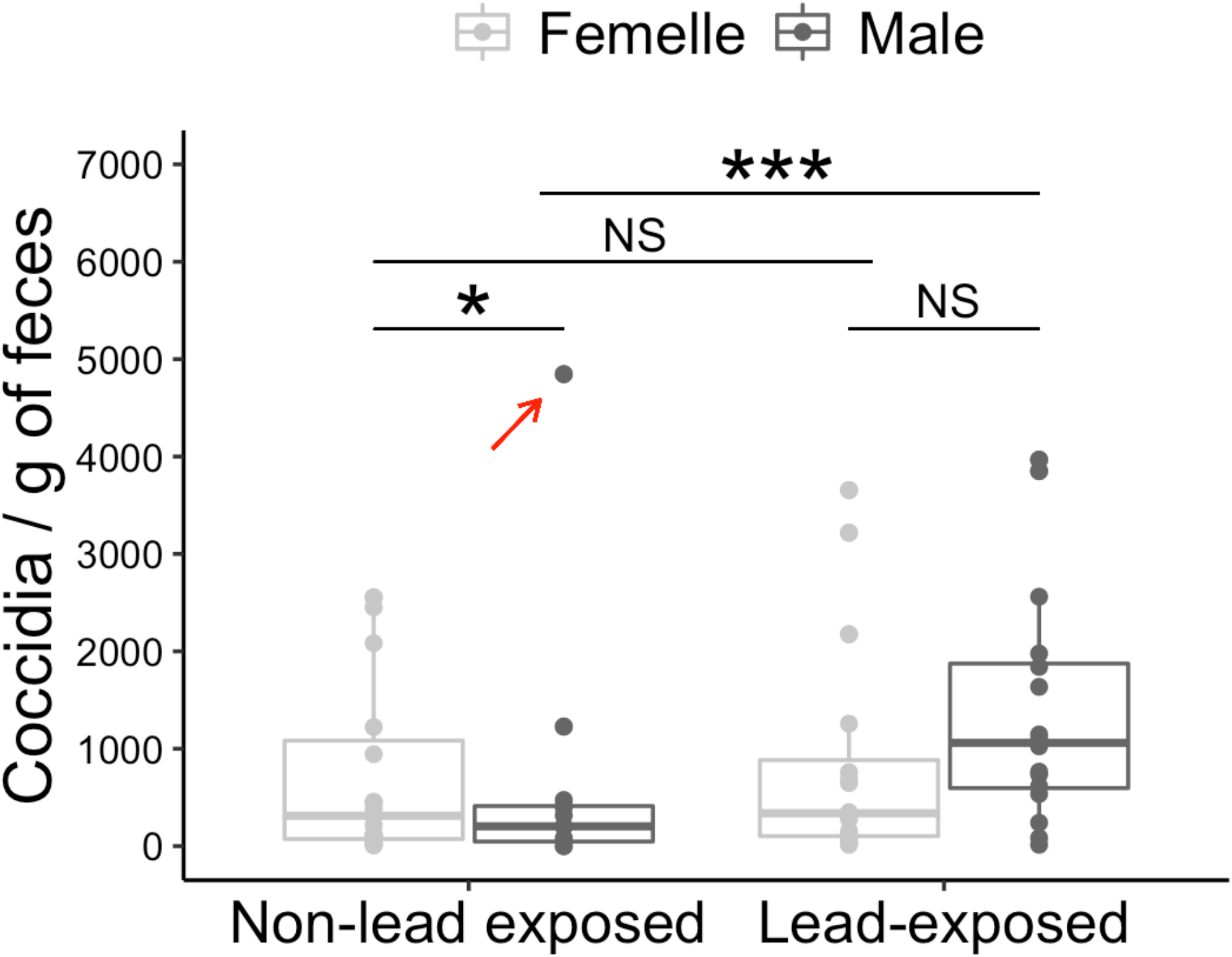
Abundance of coccidia (mean + SE) in birds non-exposed or exposed to lead among females (light grey bars) or males (dark grey bars). The red arrow indicates the extreme point in the non-lead-exposed female group, that we removed from our dataset. Significant difference between groups is indicated by an asterisk (NS: p > 0.05, *: p< 0.05, ***: p<0.001). SE, standard error.

## Results

### Lice abundance and preening behavior

We did not detect any effects of lead on preening activity (χ_1_= 0.81, P=0.37, table 1) with an average activity of 28.5±1.3 preening during 165 minutes of observation. Although we observed the presence of lice (ectoparasite abundance, for an average of 13.33 ectoparasites) in pigeons during our experiment, we did not detect any effect of lead-exposure (χ_1_= 1.24, P=0.27, table 1), time (weeks 10 and x, χ_1_= 0.84, P=0.36, table 1), or the interaction between lead exposure and time (χ_1_= 1.46, P=0.23, table 1) on ectoparasite abundance.

**Table 1.**
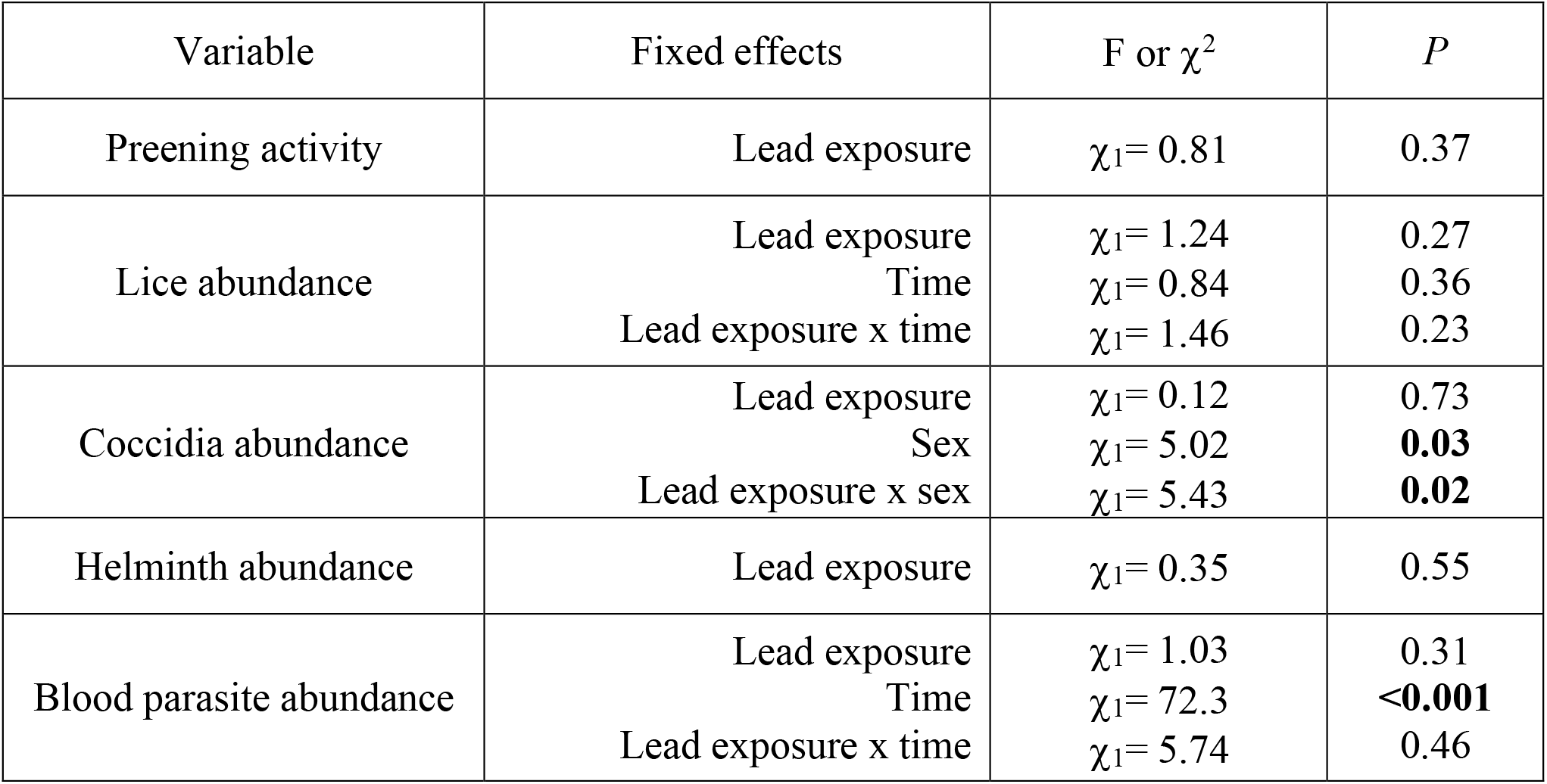
Results of models used to test the effects lead exposure on preening activity, lice abundance, coccidia abundance, helminth abundance and blood parasite abundance. Significant p-values (<0.05) are in bold in the table.

### Coccidia abundance

We detected a significant effect of the interaction between lead exposure and sex on the abundance of coccidia (mesoparasites, lead x sex: χ_1_= 5.43, P=0.02, table 1), with a higher abundance of coccidia in lead-exposed pigeons compared to the control, only in males (males: χ_1_= 12.7, P<0.001; females: χ_1_= 0.11, P= 0.74). Note that this interaction is detected only when we removed an extreme value of our dataset (see statistics section and Fig. 1). Without removing this point, we did not detect any effect of lead exposure, the sex or an interaction between lead exposure and the sex (lead x sex: χ_1_= 1.08, P=0.30, lead: χ_1_= 1.86, P=0.17, sex: χ_1_= 0.13, P=0.72)

### Helminth abundance

We did not detect any effects of lead exposure on helminth abundance (Ξ_1_= 0.35, P=0.55, table 1).

### Blood parasites abundance

The endoparasite abundance was significantly affected by time (Ξ_1_= 72.3, P<0.001, Fig. 2, table 1) but was not significantly affected by lead exposure alone (Ξ_1_= 1.03, P=0.31) or in interaction with time (Ξ_1_= 5.74, P=0.46, table 1).

**Figure 2.**
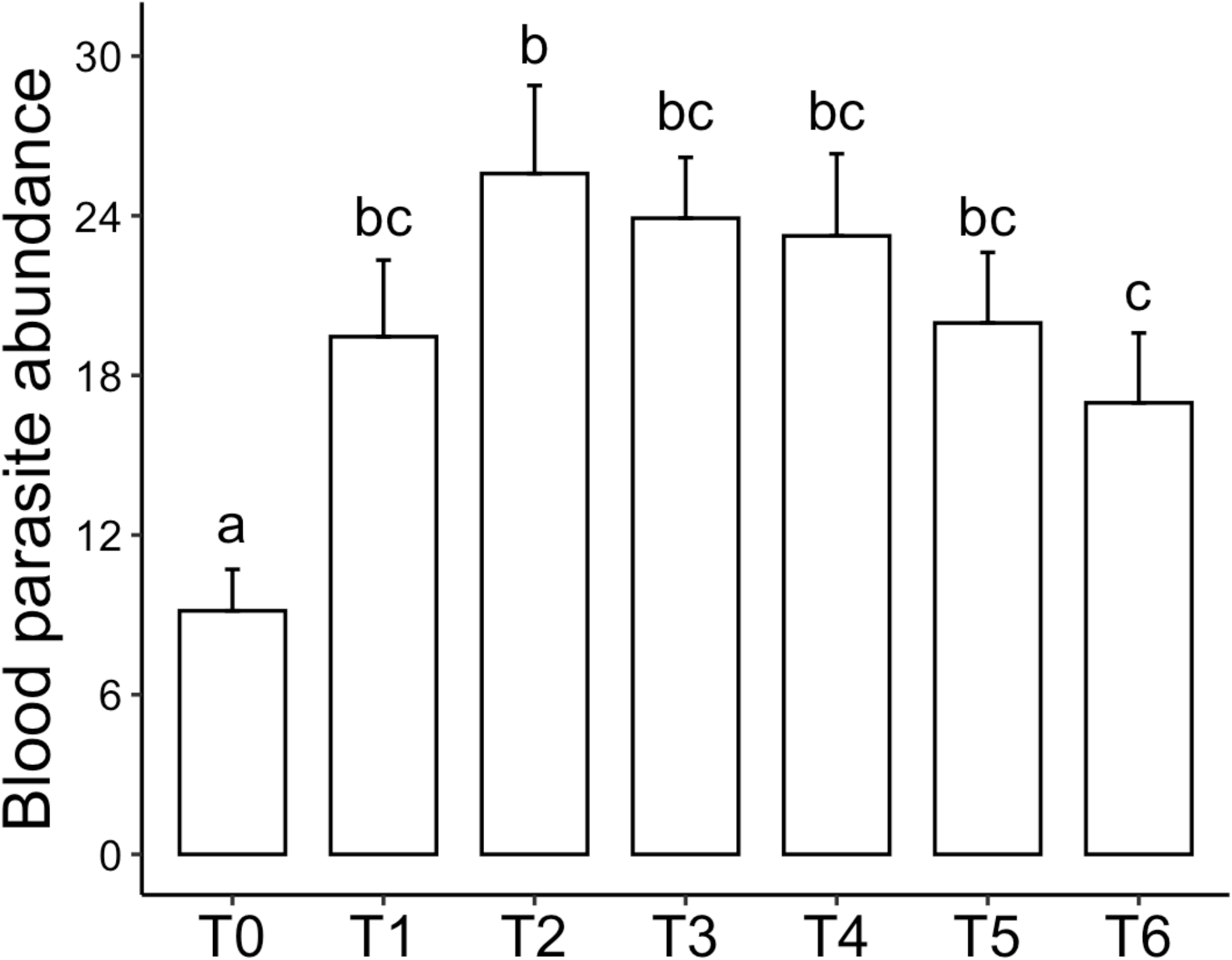
Blood parasite abundance (mean + SE) in birds among time, just before the start of the experiment (T0) and at monthly intervals thereafter (T1, T2, T3, T4, T5, T6). Significant difference between groups is indicated by different letters. SE, standard error.

## Discussion

In this study, we investigated the effect of pigeon lead exposure on their parasites with the expectation that, as we had previously shown in pigeon that lead is toxic on the humoral (Chatelain et al. 2016) and cellular immune responses (Jeantet et al., 2024), it should be profitable for parasites. In agreement with this prediction, we found a positive effect of lead exposure on coccidia abundance but only in males. In contrast, the lead exposure did not affect lice abundance, helminth abundance and blood parasites of pigeons. For ectoparasites, as the main anti-ectoparasite strategy is the preening, it is consistent with the results found on grooming behaviors (Villa et al., 2016). According to our hypothesis, the increase of parasites could be favored by host lead exposure because it impairs antiparasitic strategy and, therefore, hosts become less hostile for parasites. As the lead exposure did not impact the preening activity in our experiment (Table 1), it may explain why we did not see any effect on host lead exposure on lice abundance. Furthermore, our results are similar to those observed by Sureiro et al. (2017), who found that Patagonian rockfish (*Sebastes oculatus)* infestation by ectoparasites was similar at sites exposed to anthropogenic pollution and at non-exposed sites. They hypothesized that, due to their lifestyle outside the host and their constant exposure to environmental conditions, ectoparasites may have developed greater resistance to environmental changes.

However, this explanation does not prevail for blood parasites, heminth and coccidia abundances, for which humoral and cellular immune responses constitute the main antiparasitic strategy for the host against these meso- and endoparasites.

This is particularly the case for blood parasites, as a previous correlative study in Parisian pigeons reported a positive relationship between lead exposure and blood parasites intensity (Gasparini et al., 2014). In the present experimental study, we observed an increase over the time of blood parasites in the two treatments (control and lead-exposed, Figure 2). We suspect a seasonal effect to explain this temporal increase linked to the presence of the vector. Blood parasite abundances we measured in our study concerns three genus *Haemoproteus* spp., *Plasmodium* spp. and *Leucocytozoon* spp. In particular, Plasmodium is transmitted by mosquitoes of genus *Culex* and *Ochlerotatus* (Ferraguti et al., 2013). These plasmodium vectors have been shown to be active mainly in the summer season (Ferraguti et al., 2013). Consequently, prevalence of infected mosquitoes by plasmodium are higher in summer compared to winter and spring seasons (Ferraguti et al., 2013). These seasonal dynamics of the mosquito vector activity likely explain the increase observed during our experiment, which started in February (t0) and ended in August (t6). This strong seasonal effect on blood parasites prevalence may have hidden the effect of our lead treatment in our experiment.

Moreover, several studies have observed no correlation between immune system components and the infestation status of hosts (Rohlenová et al., 2011; Sueiro et al., 2017), which could explain why, contrary to our prediction, and despite the toxic effects of lead on the immune system of pigeons, we did not observe any effects of lead on the abundance of blood parasites and helminths. Additionally, Blanar et al. (2009) suggested that parasites indirectly exposed to pollutants, such as meso- and endoparasites, might be less vulnerable to pollutants because they are ‘protected’ by the host’s homeostatic and detoxification mechanisms, which could explain our findings. Finally, we believe that the quantification of helminth abundance performed through indirect measures (via coproscopy) might not accurately reflect their true abundance, thereby potentially masking any effects of lead exposure on these parasites.

We observed a positive effect of lead exposure on coccidian parasites, which was dependent on sex. Notably, this effect was only detected in males, in contrast to other studies on bird species that found no correlation between coccidia abundance and sex (Brown et al., 2010; Pereira et al., 2013). Although the differences we observed between males and females are consistent with previous research on pigeons, where physiological and behavioral differences between sexes, related to antiparasitic strategies, have been documented (Jeantet et al., 2024), we expected a stronger effect in females. Indeed, our previous findings indicating that females exhibit a weaker immune response compared to males (Jeantet et al., 2024). Therefore, we suggest that the increased coccidia abundance in males in lead exposure condition may be related to other physiological or behavioral factors that were not measured in this study. In contrast, without lead exposure, coccidia abundance was higher in females than in males, consistent with our earlier findings of a stronger immune response in males.

The alternative hypothesis we also tested in our study is that host lead exposure could also expose parasites to the pollutant. In this context, we expected to observe toxic effects on parasites and lower abundances in the lead-exposed group. This was not the case for the three kinds of parasites quantified. These results are inconsistent with those observed by El-Bouhy et al. (2016), who observed a negative effect of copper and lead exposure on the vitality and intensity of *Oreochromis niloticus* ectoparasites. Also, contrary to our results, Lafferty, (1997) showed negative effects of pollutants on mesoparasites. Therefore, the toxic effect of host lead exposure on parasites may depend on the specific host-parasite interaction and pollutants considered. For example, Eeva & Klemola (2013) showed that environmental pollution decreased the prevalence of the ectoparasite *Protocalliphora* sp. but has no effect on another ectoparasite, *Ornithomyia* sp. in bird *Ficedula hypoleuca*. Similarly, Sueiro et al. (2017) reported negative, positive or no effects of pollutants on several taxa of fish parasites. When parasites are more sensitive to pollutants than their hosts, their abundance would be reduced in the presence of pollutants. In contrast, when parasites are more resistant to pollutants than their hosts, their abundance would increase (El-Bouhy et al., 2016).

In conclusion, our study shows that in feral pigeons the effect of lead exposure is not toxic for their three main types of parasites: lice and coccidia (ectoparasites), helminths (mesoparasites) and blood parasites (endoparasites). In contrast, it can be beneficial to coccidia abundances, especially in males. However, it is not possible to generalize this conclusion for other systems as previous studies reported contrasting effects (negative, positive or no effect) of host pollutant exposure on their parasites. However, our study underlines the importance to consider the host environment, especially pollutants, to understand the dynamics of parasites.

## Acknowledgements

This work was supported by the French National program EC2CO (Ecosphère Continentale et Côtière). AJ’ PhD grant was funded by by the ‘Biodiversity, Evolution, Ecology, Society’ initiative of the Sorbonne University Alliance. This study was carried out in strict accordance with the recommendations of the European Convention for the Protection of Vertebrate Animals used for Experimental and Other Scientific Purposes (revised Appendix A). All experiments and captures were approved by Charles Darwin Animal Experimentation Ethics Committee and French authorities (the “Ministère de l’éducation nationale, de l’enseignement supérieur et de la recherche”, permit N°#17554 2018111610466351 ; and Ville de Paris).

## Author contributions

Aurélie Jeantet, Julien Gasparini and Fabienne Audebert conceived the ideas and designed the methodology; Aurélie Jeantet analyzed the data and led the writing of the manuscript together with Julien Gasparini, David Rozen-Rechels and Fabienne Audebert. All the authors collected the data, contributed critically to the drafts and gave final approval for publication.

## Data availability statement

The data will be deposited in Zenodo or other equivalent archives when accepted.

## References

Bagge, A. M., & Valtonen, E. T. (1996). Experimental study on the influence of paper and pulp mill effluent on the gill parasite communities of roach (Rutilus rutilus). Parasitology, 112(5), 499–508. 10.1017/S0031182000076964

Blanar, C. A., Munkittrick, K. R., Houlahan, J., MacLatchy, D. L., & Marcogliese, D. J. (2009). Pollution and parasitism in aquatic animals: A meta-analysis of effect size. Aquatic Toxicology, 93(1), 18–28. 10.1016/j.aquatox.2009.03.002

Boyce, N. P., & Yamada, S. B. (1977). Effects of a Parasite, Eubothrium salvelini (Cestoda: Pseudophyllidea), on the Resistance of Juvenile Sockeye Salmon, Oncorhynchus nerka, to Zinc. Journal of the Fisheries Research Board of Canada, 34(5), 706–709. 10.1139/f77-110

Brown, M. A., Ball, S. J., & Snow, K. R. (2010). Coccidian parasites of British wild birds. Journal of Natural History, 44(43–44), 2669–2691. 10.1080/00222933.2010.501531

Chatelain, M., Gasparini, J., & Frantz, A. (2016a). Do trace metals select for darker birds in urban areas? An experimental exposure to lead and zinc. Global Change Biology, 22(7), 2380– 2391. 10.1111/gcb.13170

Chatelain, M., Gasparini, J., & Frantz, A. (2016b). Trace metals, melanin-based pigmentation and their interaction influence immune parameters in feral pigeons (Columba livia). Ecotoxicology, 25(3), 521–529. 10.1007/s10646-016-1610-5

Chatelain, M., Gasparini, J., Haussy, C., & Frantz, A. (2016). Trace Metals Affect Early Maternal Transfer of Immune Components in the Feral Pigeon. Physiological and Biochemical Zoology, 89(3), 206–212. 10.1086/685511

Eeva, T., & Klemola, T. (2013). Variation in prevalence and intensity of two avian ectoparasites in a polluted area. Parasitology, 140(11), 1384–1393. 10.1017/S0031182013000796

Eeva, T., Ryömä, M., & Riihimäki, J. (2005). Pollution-related changes in diets of two insectivorous passerines. Oecologia, 145(4), 629–639. 10.1007/s00442-005-0145-x

El-Bouhy, Z., Reda, R., & El-Azony, A. (2016). Effect of Copper and Lead as Water Pollutants on Ectoparasitic Infested Oreochromis niloticus. Zagazig Veterinary Journal, 44(2), 156–166. 10.21608/zvjz.2016.7858

Ferraguti, M., Martínez-de La Puente, J., Muñoz, J., Roiz, D., Ruiz, S., Soriguer, R., & Figuerola, J. (2013). Avian Plasmodium in Culex and Ochlerotatus Mosquitoes from Southern Spain: Effects of Season and Host-Feeding Source on Parasite Dynamics. PLoS ONE, 8(6), e66237. 10.1371/journal.pone.0066237

Gasparini, J., Jacquin, L., Laroucau, K., Vorimore, F., Aubry, E., Castrec-Rouëlle, M., & Frantz, A. (2014). Relationships Between Metals Exposure and Epidemiological Parameters of Two Pathogens in Urban Pigeons. Bulletin of Environmental Contamination and Toxicology, 92(2), 208–212. 10.1007/s00128-013-1172-7

Jacquin, L., Lenouvel, P., Haussy, C., Ducatez, S., & Gasparini, J. (2011). Melanin-based coloration is related to parasite intensity and cellular immune response in an urban free living bird: The feral pigeon Columba livia. Journal of Avian Biology, 42(1), 11–15. 10.1111/j.1600-048X.2010.05120.x

Jeantet, A., Sandmeyer, L., Campech, C., Audebert, F., Agostini, S., Pellerin, A., & Gasparini, J. (2023). The “parasite detoxification hypothesis”: Lead exposure potentially changes the ecological interaction from parasitism to mutualism. Ecotoxicology, 32(5), 666–673. 10.1007/s10646-023-02678-z

Jeantet, A., Audebert, F., Agostini, S., Decencière, B., Lorang, C., Jamet, L., Giner, C., Flandi, E., Nardou, N., Lemaire, B., Federici, P., Rozen-Rechels, D., & Gasparini, J. (2024). Crossed effects of helminth infection and lead exposure on fitness: An experimental study in feral pigeons (Columba livia). Journal of Animal Ecology, 1365-2656.14211. 10.1111/1365-2656.14211

Koop, J. A. H., & Clayton, D. H. (2013). Evaluation of two methods for quantifying passeriform lice: Quantifying Ectoparasites on Passerines. Journal of Field Ornithology, 84(2), 210–215. 10.1111/jofo.12020

Koop, J. A. H., Huber, S. K., Laverty, S. M., & Clayton, D. H. (2011). Experimental Demonstration of the Fitness Consequences of an Introduced Parasite of Darwin’s Finches. PLoS ONE, 6(5), e19706. 10.1371/journal.pone.0019706

Korine, C., Pilosof, S., Gross, A., Morales-Malacara, J. B., & Krasnov, B. R. (2017). The effect of water contamination and host-related factors on ectoparasite load in an insectivorous bat. Parasitology Research, 116(9), 2517–2526. 10.1007/s00436-017-5561-4

Lafferty, K. D. (1997). Environmental parasitology: What can parasites tell us about human impacts on the environment? Parasitology Today, 13(7), 251–255. 10.1016/S0169-4758(97)01072-7

Lefcort, H., Aguon, M. Q., Bond, K. A., Chapman, K. R., Chaquette, R., Clark, J., Kornachuk, P., Lang, B. Z., & Martin, J. C. (2002). Indirect Effects of Heavy Metals on Parasites May Cause Shifts in Snail Species Compositions. Archives of Environmental Contamination and Toxicology, 43(1), 34–41. 10.1007/s00244-002-1173-8

Lewis, J., Hoole, D., & Chappell, L. H. (2003). Parasitism and environmental pollution: Parasites and hosts as indicators of water quality. Parasitology, 126(7), S1–S3. 10.1017/S0031182003003962

Marcogliese, D. (2008). The impact of climate change on the parasites and infectious diseases of aquatic animals. Revue Scientifique et Technique (International Office of Epizootics), 27, 467–484.

Pascoe, D., & Cram, P. (1977). The effect of parasitism on the toxicity of cadmium to the three-spined stickleback, Gasterosteus aculeatus L. Journal of Fish Biology, 10(5), 467–472. 10.1111/j.1095-8649.1977.tb04079.x

Pereira, L. Q., Corrêa, I. M. O., Schneiders, G. H., Linhares, M. T., Almeida, D. T., & Lovato, M. (2013). Isospora bocamontensis (Protozoa: Apicomplexa) in captive yellow cardinal Gubernatrix cristata (Passeriformes: Emberezidae). Pesquisa Veterinária Brasileira, 33(3), 384– 388. 10.1590/S0100-736×2013000300018

Price, P. W. (1980). PAR volume 81 issue 3 Cover and Back matter. Parasitology, 81(3), b1– b11. 10.1017/S0031182000061850

Raynaud, J.-P., William, G., & Brunault, G. (1970). Etude de l’efficacité d’une technique de coproscopie quantitative pour le diagnostic de routine et le contrôle des infestations parasitaires des bovins, ovins, équins et porcins. Annales de Parasitologie Humaine et Comparée, 45(3), 321–342. 10.1051/parasite/1970453321

Rohlenová, K., Morand, S., Hyršl, P., Tolarová, S., Flajšhans, M., & Šimková, A. (2011). Are fish immune systems really affected by parasites? An immunoecological study of common carp (Cyprinus carpio). Parasites & Vectors, 4(1), 120. 10.1186/1756-3305-4-120

Sanchez-Ramirez, C., Vidal-Martinez, V. M., Aguirre-Macedo, M. L., Rodriguez-Canul, R. P., Gold-Bouchot, G., & Sures, B. (2007). CICHLIDOGYRUS SCLEROSUS (MONOGENEA: ANCYROCEPHALINAE) AND ITS HOST, THE NILE TILAPIA (OREOCHROMIS NILOTICUS), AS BIOINDICATORS OF CHEMICAL POLLUTION. Journal of Parasitology, 93(5), 1097–1106. 10.1645/GE-1162R.1

Sueiro, M. C., Bagnato, E., & Palacios, M. G. (2017). Parasite infection and immune and health-state in wild fish exposed to marine pollution. Marine Pollution Bulletin, 119(1), 320–324. 10.1016/j.marpolbul.2017.04.011

Villa, S. M., Goodman, G. B., Ruff, J. S., & Clayton, D. H. (2016). Does allopreening control avian ectoparasites? Biology Letters, 12(7), 20160362. 10.1098/rsbl.2016.0362

Wobeser, G. A. (2008). Parasitism: Costs and Effects. In C. T. Atkinson, N. J. Thomas, & D. B. Hunter (Eds.), Parasitic Diseases of Wild Birds (1st ed., pp. 1–9). Wiley. 10.1002/9780813804620.ch1

